# Adverse PFAS effects on mouse oocyte *in vitro* maturation are associated with carbon-chain length and inclusion of a sulfonate group

**DOI:** 10.1101/2022.05.30.493919

**Authors:** Jianan Feng, Edgar J. Soto-Moreno, Aashna Prakash, Ahmed Z. Balboula, Huanyu Qiao

## Abstract

Per- and polyfluoroalkyl substances (PFAS) are man-made chemicals that are used in products such as non-stick cookware, stain-resistant coating, and food packaging. PFAS are characterized by their fluorinated carbon chains that make them hard to degrade and bioaccumulate in human and animals. Toxicological studies have shown PFAS toxic effects: cytotoxicity, immunotoxicity, neurotoxicity, and reproductive toxicity. Two major categories of PFAS are perfluoroalkyl carboxylic acid (PFCA) and perfluoroalkyl sulfonic acid (PFSA). In this study, we used a mouse-oocyte-*in*-*vitro*-maturation (IVM) system to study how the structures of PFAS, such as carbon-chain length and functional groups, determine their reproductive toxicity. We found the toxicity of PFAS is elevated with increasing carbon-chain length and the inclusion of the sulfonate group. Specifically, at 600 µM, perfluorohexanesulfonic acid (PFHxS) and perfluorooctanesulfonic acid (PFOS) reduced the rates of both germinal vesicle breakdown (GVBD) and polar body extrusion (PBE) as well as induced the formation of relatively large polar bodies. However, the shorter PFSA, perfluorobutanesulfonic acid (PFBS), and all PFCA did not show similar adverse cytotoxicity. We further examined mitochondria and cytoskeleton, two essential factors for cell division, in PFOS- and PFHxS-treated oocytes. We found that 600 µM PFHxS and PFOS exposure induced excess reactive oxygen species (ROS) and decreased mitochondrial membrane potential (MMP). Cytoskeleton analysis revealed that PFHxS and PFOS exposure induced chromosome misalignment, abnormal F-actin organization, elongated the spindle formation, and symmetric division in the treated oocytes. Together, our study provides new information on the structure-toxicity relationship of PFAS.

**Synopsis:** Reproductive toxicity of PFAS, a group of persistent organic pollutants, is determined by their chemical structures.

## 1. Introduction

Since the start of their production in the 1940s, per- and polyfluoroalkyl substances (PFAS) have been employed as surfactants and polymers^1^ in various industry branches including aerospace, biotechnology, and mining.^2^ Carbon-fluorine bonds in PFAS make them extremely stable and resistant to degradation.^3^ Such properties, on the other hand, cause PFAS to accumulate in animal bodies as persistent organic pollutants.^4^ Human are regularly exposed to PFAS through inhalation, dermal exposure, food, and drinking water.^5^ Two subcategories of PFAS (Table 1), perfluoroalkyl carboxylic acids (PFCA) and perfluoroalkyl sulfonic acids (PFSA) have drawn great attention in recent years due to their demonstrated neurotoxicity^6,7^, developmental toxicity^8^, immunotoxicity^9^, hepatotoxicity^10^, and especially reproductive toxicity^11,12^. Unfortunately, both PFCA and PFSA have been detected in humans. For instance, one of the PFSA, perfluorooctanesulfonic acid (PFOS), has a median serum concentration as high as 12.70 ng/mL.^13^ In terms of female reproductive toxicity, PFCA and PFSA have been shown to be able to pass through the blood-follicle barrier and can be detected in follicular fluid.^14^ Clinical evidence shows that PFCA and PFSA are associated with a late age at menarche, irregular menstrual cyclicity, and early menopause.^14^ *In vitro* studies have demonstrated the direct cytotoxicity of PFCA and PFSA on mouse oocyte maturation. Oocyte maturation releases oocytes from dictyate arrest and prepares them for fertilization. Dramatic morphological changes occur during this process, including germinal vesicle breakdown (GVBD) and polar-body extrusion (PBE). Various epigenetic regulations are known to be involved in oocyte maturation, including histone acetylation, phosphorylation, and SUMOylation.^15^ GVBD, the breakdown of nuclear envelop, exposes the chromosomes to many environmental toxicants, such as iodoacetic acid (IAA)^16^, PM ^17^ and several PFAS chemicals^18–21^. IAA, for example, has been shown to induce DNA damage and cause chromosome misalignment at the metaphase I stage.^16^ PFOS exposure has also been shown to alter histone methylation levels with increased H3K4me3 and decreased H3K9me3^19^ and, furthermore, modulate maternal-to-zygotic transition^22,23^. Mitochondria also play important roles during oocyte maturation because large amounts of ATP are required for continuous transcription and translation.^24^ Mitochondrial DNA (mtDNA) in oocytes can encode various functional proteins including ATP synthase, cytochrome oxidase, NADH, and pan-reductase. Therefore, mitochondrial dysfunction caused by PFAS can result in excess ROS generation, dissipation of the mitochondrial membrane potential, and early apoptosis. These mitochondria-related problems have been found in oocytes^18,19^ and other cell types ^25–27^.

**Table 1.** PFAS chemicals used in this study. PFBA, PFHxA, and PFOA are categorized as perfluoroalkyl carboxylic acid (PFCA), whereas PFBS, PFHxS, and PFOS are perfluoroalkyl sulfonic acid (PFSA). The number of carbons in each chemical is listed for reference.

| PFAS | Full name | Structure | Categories | #Carbon |
| --- | --- | --- | --- | --- |
| PFBA | perfluorobutanoic acid | $\text{CF}_3(\text{CF}_2)_2\text{COOH}$ | PFCA | 4 |
| PFHxA | perfluorohexanoic acid | $\text{CF}_3(\text{CF}_2)_4\text{COOH}$ | PFCA | 6 |
| PFOA | perfluorooctanoic acid | $\text{CF}_3(\text{CF}_2)_6\text{COOH}$ | PFCA | 8 |
| PFBS | potassium nonafluorobutanesulfonate | $\text{CF}_3(\text{CF}_2)_3\text{KO}_3\text{S}$ | PFSA | 4 |
| PFHxS | potassium perfluorohexanesulfonate | $\text{CF}_3(\text{CF}_2)_5\text{KO}_3\text{S}$ | PFSA | 6 |
| PFOS | potassium perfluorooctanesulfonate | $\text{CF}_3(\text{CF}_2)_7\text{KO}_3\text{S}$ | PFSA | 8 |

Recently, the cytotoxicity of individual long-chain PFAS on mouse oocytes has been widely studied. However, the data on the cytotoxicity of short-chain PFAS, such as PFBA and PFBS, are still missing. Furthermore, various experimental conditions in different studies make it hard to distill the data. In the present study, we used a mouse-oocyte-*in-vitro*-maturation model to systematically compare the toxicity of six PFAS and to elucidate which factors determine the toxicity of PFAS on oocyte maturation. We found that PFSA is more toxic than PFCA, and that toxicity is positively correlated to carbon-chain length. Interestingly, by calculating the size ratio between the first polar body (PB) and the oocyte, we noticed large PBs occurred in the PFSA group, but not in the PFCA group. To study how 600 µM PFHxS and PFOS disrupted meiosis, we analyzed the mitochondrial functions and cytoskeleton structures of mouse oocytes. We found a significant increase in ROS levels, chromosome misalignment, and frequency of abnormally elongated spindles in PFHxS/PFOS-treated oocytes compared to the untreated controls. Moreover, the mitochondrial membrane potential decreased in treated oocytes compared to untreated controls. We also demonstrated that aberrant F-actin distribution can cause the large PB phenotype observed in the PFHxS and PFOS treated groups. Collectively, our data provide new evidence for the structure-toxicity relationship of PFAS.

## 2. Methods and Materials

### 2.1. Chemicals

All PFAS chemicals listed in Table 1 are from Synquest Laboratories (Alachua, FL, USA). Dimethyl sulfoxide (DMSO) was sourced from Avantor (Allentown, PA, USA). All other chemicals were purchased from Sigma-Aldrich (St. Louis, MO, USA).

### 2.2. Animals

In this study, we used two mouse strains, CD-1 (Charles River Laboratories, Wilmington, MA) and CF-1 (Envigo, Indianapolis, Indiana) to confirm our results. They are housed in the Animal Care Facility at the University of Illinois Urbana-Champaign (UIUC) and the University of Missouri-Columbia, respectively. Mice were housed under 12 h dark/12 h light cycles at 22 ± 1 °C and were provided food and water ad libitum. Animal handling and procedures were approved by the UIUC Institutional Animal Care and Use Committee and the University of Missouri Animal Care and Use Committee.

### 2.3. Mouse oocyte *in vitro* maturation

Female mice (4-6 weeks old) were euthanized and dissected for ovary collection. The ovaries were washed in pre-warmed M2 media, which contained 100 μM IBMX, before the isolation of cumulus-oocyte-complexes (COCs). This was accomplished by using sterile syringe needles to break down the antral ovarian follicles. Cumulus cells then were removed through repeated pipetting. Viable denuded oocytes were collected in pre-warmed M2 media with 100 μM IBMX. During the oocyte collection process, IBMX arrests oocytes at prophase I by inhibiting adenylate cyclase and elevating cAMP levels to hinder oocyte nuclear maturation.^28^ Oocytes were washed in and transferred to pre-warmed M16 media covered by mineral oil. The oocytes were incubated at 37°C in a 5% CO_2_ incubator and examined at 2 h and 14 h – the stages at which oocytes typically reach GVBD and already extruded the PB, respectively.

### 2.4. Chemical treatment

Each PFAS (PFOA, PFHxA, PFBA, PFOS, PFHxS, and PFBS) was dissolved in DMSO and diluted to a final concentration of 600 μM in M16 media because PFOA and PFOS have been proven to compromise mouse oocyte *in vitro* maturation at this concentration.^18,19^. The same amount of DMSO was also added into M16 media as a vehicle control. The amount of DMSO did not exceed 0.1% in all cases.

### 2.5. Calculating the size ratio between the first PB and the oocyte

After 14 h culture at 37°C in a 5% CO_2_ incubator, oocytes were imaged under a Nikon A1R confocal microscope. The cross-section areas of both first PB and oocyte were measured using either NIS-Elements software or Fiji software^29^, and the area ratio between them was calculated to detect abnormal oocyte division.

### 2.6. Measurement of mitochondrial membrane potential (MMP)

MMP was measured by using positively charged tetramethylrhodamine, methyl ester (TMRM) as a probe. Briefly, oocytes were incubated at 37°C in a 5% CO_2_ incubator for 8 h before being washed with M2 medium 2-3 times to remove PFAS chemicals in the treatment groups. Next, the oocytes were incubated in M2 media containing 25 nM TMRM at 37°C for 30 min in the dark. The stained oocytes were washed 2-3 times in M2 media to remove extra TMRM. The orange signal from the polarized mitochondria was immediately examined under an Olympus IX73 microscope with a consistent parameter setting. The red signal intensity of the ROI (region of interest) was quantified using Fiji software.

### 2.7. Measurement of intracellular reactive oxygen species (ROS) levels

2’,7’-dichlorofluorescin diacetate (DCFH-DA), a fluorescent probe sensitive to oxidation, was used to measure the intracellular ROS level in oocytes. Treated and untreated oocytes were cultured for 8 h at 37°C in a 5% CO_2_ incubator. PFAS chemicals were removed by washing oocytes in M2 media 2-3 times. Next, oocytes were transferred to M2 media with 5 µM DCFH-DA for another 30 min incubation. Additional DCFH-DA was washed off with fresh M2 media. The green signal was captured under an Olympus IX73 microscope with the same imaging parameters for control and all the experimental groups; the intensity of signals from the ROI was analyzed using Fiji software.

### 2.8. Immunocytochemistry and fluorescence microscopy

Oocytes were fixed for 20 minutes in 1× phosphate buffer saline (PBS) with 3.7% paraformaldehyde (MilliporeSigma P6148) at room temperature. Fixed oocytes were permeabilized in PBS with 0.1% Triton X-100 for 20 minutes at room temperature, followed by a 20-minute incubation in blocking solution (0.3% BSA and 0.01% Tween-20 dissolved in PBS). Primary antibody incubation was performed at room temperature for 1 hour. Oocytes were then washed three times (7 minutes each) in blocking solution. To detect F-actin and the meiotic spindle, Texas Red X Phalloidin (1:50, ThermoFisher Scientific T7471) and anti-rabbit α-tubulin monoclonal antibody Alexa® 488 conjugate (1:100, Cell Signaling, 5063S) were used, respectively. Oocytes were mounted on slides using VECTASHIELD with 4’,6-Diamidino-2-Phenylindole, Dihydrochloride (DAPI; Invitrogen 2116137). Fluorescence signals were detected under a 100× immersion oil objective using Leica TCS SP8 diode confocal microscope. Z-plane images were captured to span the entire oocyte at 2 µm Z-intervals.

### 2.9. Time-lapse confocal microscopy

Germinal vesicle oocytes were cultured in pre-warmed and equilibrated maturation medium and imaged over time under a 40× immersion oil objective using a Leica TCS SP8 confocal microscope equipped with a microenvironmental chamber to regulate the temperature and CO_2_ at 37 °C and 5% in humidified air. SiR-tubulin (Cytoskeleton NC0958386) was added to the maturation medium to label microtubules.^30,31^ Bright-field and 647 nm wavelength images acquisition were started at 1.5 hours after collection (30-45 minutes collection time), in which the oocytes were at the GVBD stage. Time-lapse images were taken every 30 minutes. Z-plane images were captured to span the entire oocyte at 7 µm Z-intervals.

### 2.10. Statistical analysis

All experiments were repeated at least three times. The data were presented as mean ± SEM. One way analysis of variance (ANOVA) was used to compare means between multiple groups, followed by Tukey post hoc procedure. Unpaired two-tailed t-tests were used to compare means between two groups. All analysis was done using R (version 4.0.3, Vienna, Austria). Graphs were made using OriginPro 2020 (Northampton, MA, USA). Comparisons were considered significant at * P < 0.05, ** P < 0.01, and *** P < 0.001.

## 3. Results

### 3.1. PFAS with a long carbon chain impede mouse oocyte maturation

To determine the effect of carbon-chain length on the toxicity of PFAS, comparisons of GVBD rate, PBE rate, and relative PB size (size ratio) were conducted among PFCA and PFSA groups. We found the toxicity of PFSA is positively correlated with the carbon-chain length. Specifically, GVBD and PBE rates were normal in the PFBS- and PFHxS-treated oocytes. However, as the carbon-chain length increased to eight, PFOS treatment resulted in a lower GVBD rate (32.84 ± 7.21 %, versus 76.03 ± 1.11 % in the control, P < 0.001, Fig. 1B & C) and a lower PBE rate (24.93 ± 8.14%, compared to 73.84 ± 1.78 % in the control, P < 0.001, Fig. 1B & D). Interestingly, we noticed that although PFHxS-treated oocytes had normal GVBD and PBE rates, like PFOS, some oocytes extruded a large PB (Fig. 1B), a phenotype that has been qualitatively described in previous PFAS studies.^19,32,33^ Therefore, we calculated the average size ratio between the first PB and its oocyte for each treatment, as described in section 2.5. We found the average size ratio was 14.24 % in the control; in contrast, the ratios were 31.22 % in 600 µM PFHxS group and 52.57 % in the PFOS group, respectively (P < 0.001, Fig. 1B & E). Together, these results suggest a positive relationship between the toxicity of PFAS and the carbon-chain length.

**Figure 1.**
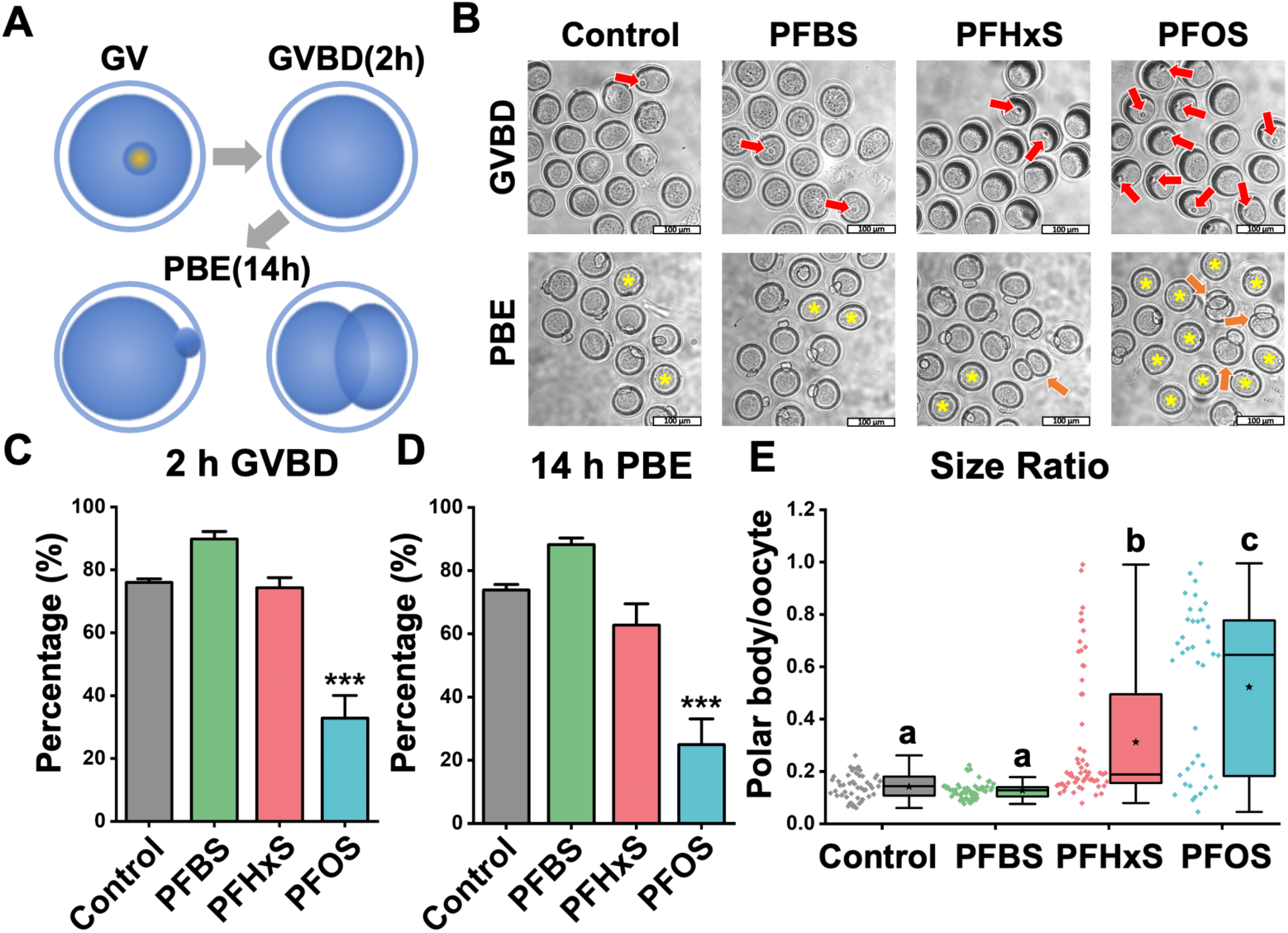
PFSA impedes mouse oocyte maturation in a carbon-chain-length-dependent manner. (A) Scheme of mouse oocyte *in vitro* maturation process. We examined GVBD at 2 h and PBE at 14 h. The bottom-right image illustrates an extrusion of an abnormal large PB. (B) Representative images show GVBD and PBE of four treatment groups (untreated, 600 µM PFBS, 600 µM PFHxS, and 600 µM PFOS). The red arrows indicate oocytes that retained their germinal vesicles after 2 hours of culture. The yellow asterisks indicate oocytes that did not extrude a PB after 14 hours of culture. Orange arrows highlight oocytes with large PBs. Scale bar: 100 µm. (C) The rates of GVBD in the control and PFSA-treated groups. (D) The rates of PBE in the control and PFSA-treated groups. A total of 136 oocytes in the control group, 116 oocytes in the PFBS-treated group, 179 oocytes in the PFHxS-treated group, and 188 oocytes in the PFOS-treated group were analyzed to calculate the GVBD and PBE rates. (E) The size ratios of PBs to oocytes in the control and PFSA-treated groups. A total of 49 oocytes in the control group, 57 oocytes in the PFBS-treated group, 63 oocytes in the PFHxS-treated group, and 40 oocytes in the PFOS-treated group were measured to calculate the size ratios. Data in bar chart were presented as mean ± SEM. All groups had at least 3 independent groups. *** P < 0.001, compared with control. Groups with different letters in the box plot have significant differences.

### 3.2. PFSA has higher and unique toxicity effects on mouse oocyte maturation compared to PFCA

Although the carbon backbones of PFAS are usually very stable due to their strong carbon-fluorine bond^34^, the functional groups, such as carboxylate and sulfonate, may have a greater influence on the toxicity of PFAS chemicals. To examine the relative toxicity of PFSA to its corresponding PFCA with the same carbon chain length (Supplemental Fig. 1), we rearranged our data in the section 3.1 to compare PFAS with the same carbon length but different functional groups (carboxylate versus sulfonate). Compared to 600 µM PFOA treatment, 600 µM PFOS-treated oocytes had a significantly lower GVBD rate (32.84 ± 7.21 % versus 74.16 ± 5.41 % in the 600 µM PFOA treatment group, P < 0.01, Fig. 2A left) and PBE rate (24.93 ± 8.14 % versus 66.50 ± 4.98 % in the 600 µM PFOA treatment group, P < 0.001, Fig. 2A right). In addition, the treatments of both 600 µM PFHxS and PFOS induced larger PBs; while the treatments of 600 µM PFHxA and PFOA could not induce larger PBs (31.22 ± 3.12 % in the PFHxS group versus 16.93 ± 2.99 % in the PFHxA group, and 52.57 ± 4.89 % in the PFOS group versus 16.84 ± 1.32 % in the PFOA group, both with P < 0.001, Fig. 2B). Based on our finding, we concluded that PFSA has a higher reproductive toxicity than PFCA with the same carbon-chain length.

**Figure 2.**
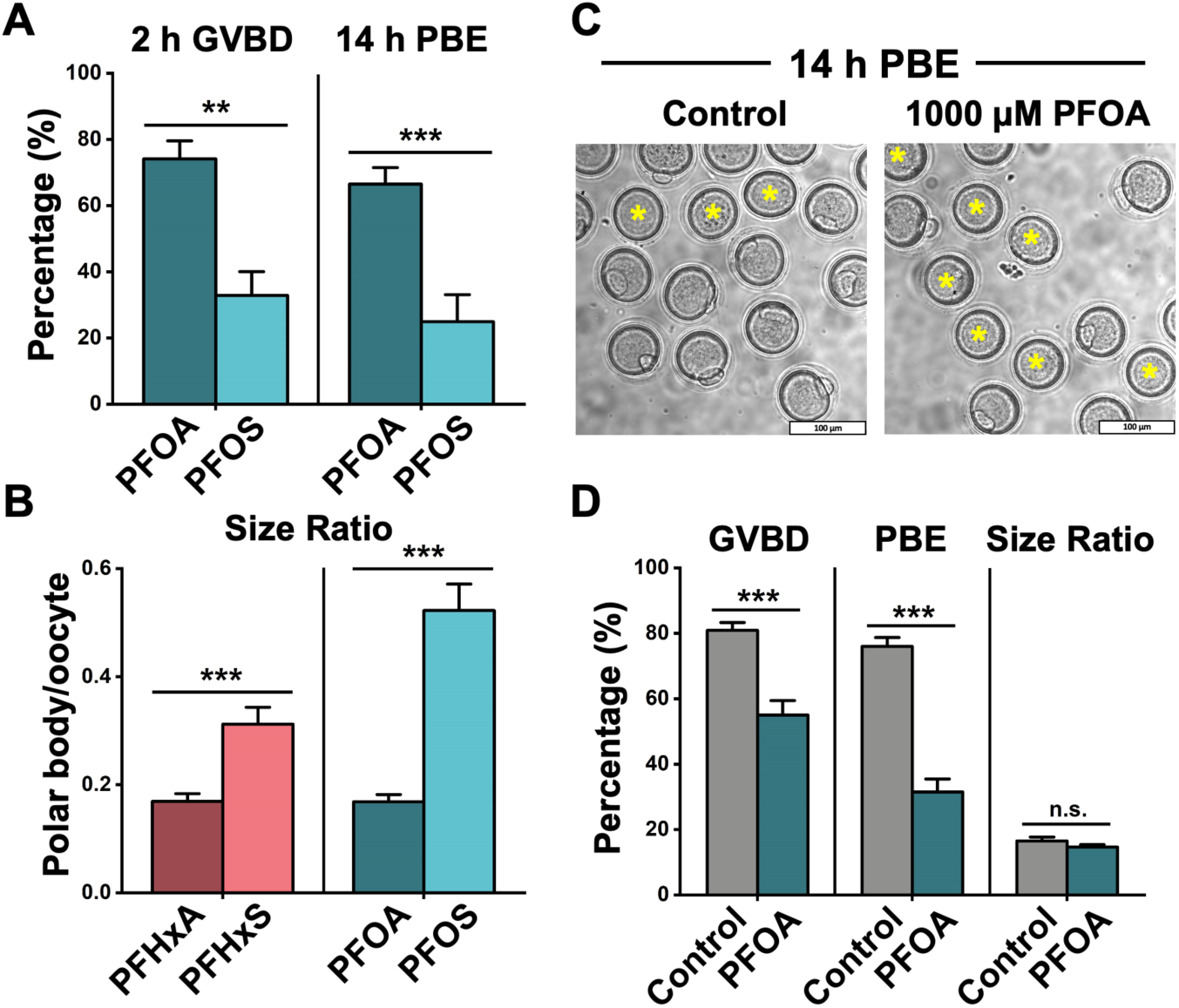
PFSA are more toxic than the PFCA with the same carbon-chain length. (A) The rates of GVBD and PBE in the PFOA- and PFOS-treated groups. The PFOA-treated group consists of 144 oocytes, and the PFOS-treated group consists of 118 oocytes. (B) The size ratios of PBs to oocytes in the PFHxA-, PFHxS-, PFOA-, and PFOS-treated groups. A total of 56 oocytes are in the PFHxA-treated group, 63 oocytes are in the PFHxS-treated group, 63 oocytes are in the PFOA-treated group, and 40 oocytes are in the PFOS-treated group. (C) Representative images showed PBE in the control and the 1000 µM PFOA-treated groups. Scale bar: 100 µm. (D) Comparison of GVBD, PBE, and size ratio between the control and 1000 µM PFOA groups. A total of 221 oocytes in the control and 346 oocytes in the 1000 µM PFOA-treated group were analyzed for GVBD and PBE rate. 40 oocytes in the control and 25 oocytes in the 1000 µM PFOA-treated group were analyzed for calculating size ratio. Data were presented as mean ± SEM. All groups had at least 3 independent groups. ** P < 0.01, *** P < 0.001, n.s.: no significance.

Next, due to the abovementioned difference, we asked whether PFSA and PFCA can disrupt oocyte maturation in different ways. We increased the concentration of PFOA to 1000 µM to induce abnormality. We found at this concentration, the GVBD and PBE rates decreased to 55 ± 4% and 31 ± 4%, but unlike PFHxS- and PFOS-treated groups, the size ratio remained within a normal range (Fig. 2C). This indicated that the large PB is a unique phenotype that is associated with PFSA, but not PFCA. To further study the specific toxic effects of PFSA, we selected 600 µM PFHxS- and PFOS-treatment groups to check the mitochondrial function and cytoskeleton structures in mouse oocytes.

### 3.3. PFHxS and PFOS elevate the level of reactive oxygen species (ROS)

For all cells that undergo aerobic respiration, redox homeostasis is maintained between the ROS generation and scavenging.^35^ Various PFAS chemicals have been reported to break the redox balance in mouse oocytes, including PFNA^20^, PFOS^19^, and PFOA^21^. Excess ROS induces oxidative stress, leading to DNA damage, protein malfunction, and lipid chain breakage.^36^ To determine if PFHxS can induce oxidative stress in oocytes and to confirm previous results for PFOS^19^, we used DCFH-DA as a probe to detect intracellular H_2_O_2_ and oxidative stress.^37^ After passively diffusing into the oocytes, DCFH-DA is cleaved by esterase to form DCFH, which can be oxidized to DCF and emit green fluorescence signal.^38^ We found PFHxS and PFOS elevated the intracellular ROS level in a carbon-chain-length-dependent manner (Fig. 3A). Our quantitative analysis (Fig. 3B) confirmed this observation: the relative fluorescent intensities were 3.09 ± 0.16 in the PFHxS-treated oocytes (versus 2.37 ± 0.16 in control, p < 0.05) and 4.19 ± 0.22 in the PFOS-treated oocytes (p < 0.001 versus control and PFHxS-treated oocytes).

**Figure 3.**
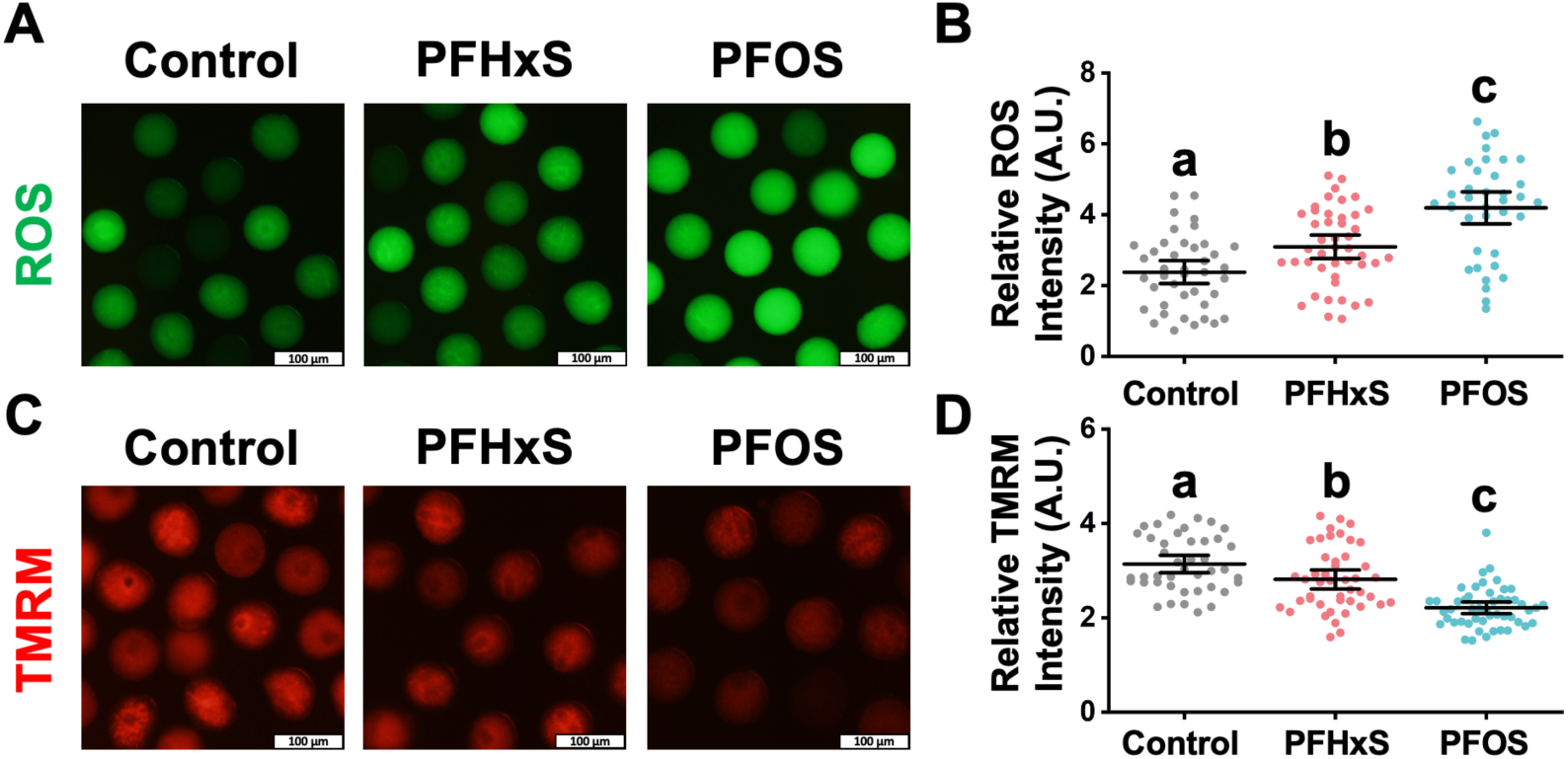
PFHxS and PFOS (600 µM) increased ROS level and reduced mitochondria membrane potential (MMP). Intracellular reactive oxygen species (ROS) levels indicated by the intensity of green DCF fluorescence signal. Quantitative analysis of the ROS levels in the control, PFHxS- and PFOS-treated groups. A total of 41 oocytes in the control, 44 oocytes in the PFHxS-treated group, and 38 oocytes in the PFOS-treated group were measured under the same camera setting. (C) MMP indicated by red TMRM intensity levels. (D) Quantitative analysis of the MMP in different treatment groups. A total of 42 oocytes in the control, 45 oocytes in the PFHxS-treated group, and 49 oocytes in the PFOS-treated group were measured. Groups with different letters in (B) and (D) are significantly different (p < 0.05). Scale bar: 100 µm.

### 3.4. PFHxS and PFOS induce mitochondrial depolarization

A positive loop exists between mitochondrion-derived ROS accumulation and mitochondrial depolarization^39^, an event that is universally associated with apoptosis and cell death.^40^ To quantify the mitochondrial membrane potential change after PFHxS and PFOS exposure, TMRM, a small cationic lipophilic fluorescent indicator^41^, was used to bind negatively charged mitochondrial membrane.^42^ Our results show that the red signal of TMRM was dimmer in PFHxS and PFOS treatment groups, again, depending on the carbon-chain lengths (Fig. 3C). Statistically, the red signal intensity is 3.14 ± 0.09 in control, while it is 2.81 ± 0.10 in the PFHxS-treatment group (p < 0.05 versus control) and 2.12 ± 0.06 in the PFOS-treatment group (p < 0.001 versus control and PFHxS group). Therefore, we concluded that 600 µM PFHxS and PFOS cause mitochondrial depolarization. Mitochondrial depolarization is regarded as a sign of early apoptosis. Using time lapse confocal microscopy, we observed nearly 30% of PFOS-treated oocytes (600 µM) underwent oocyte blebbing which could be a sign of an apoptotic event (Supplemental Fig. 2 PFOS). Interestingly, we frequently observed the fusion of the PB/cytoplasmic blebbing and the oocyte (supplemental Fig. 2) indicating cytokinesis failure.

### 3.5. PFHxS and PFOS cause chromosome misalignment and abnormal assembly of spindle and F-actin

Abnormal first PB morphology, including fragmented and enlarged PB, is associated with poor outcomes of *in vitro* fertilization (IVF).^43–45^ Therefore, the morphology of the first PB is used as a prognostic factor regarding egg quality in IVF clinics.^43,44^ The position of the spindle within the cell determines the cleavage plane. Therefore, during oocyte meiosis I, the centrally located spindle migrates towards the cortex, a necessary process to extrude a tiny PB. This mechanism ensures that the egg retains virtually all RNAs and proteins (synthesized during oogenesis) necessary for fertilization and early embryo development. Active spindle migration towards the cortex is driven by dynamic F-actin. Thus, F-actin inhibition or stabilization prevents spindle migration towards the cortex and leads to cytokinesis failure.^46-49^ Since our observation indicated that PFSA, but not PFCA, resulted in the extrusion of enlarged PB with frequent oocyte blebbing (Fig. 2B), we hypothesized that PFSA perturbs cytoskeleton organization within the oocyte. To this end, we did cytological staining of DNA/chromosome, microtubules (to label the spindle), and F-actin in oocytes from control, 600 µM PFHxS-, and 600 µM PFOS-treated groups. We found a significant increase of stabilized F-actin signal around the spindle regions in the PFHxS and PFOS treatment groups (Fig. 4A and 4B), which could hinder the migration of spindle to the oocyte cortex. In addition, the F-actin cage surrounding the spindle (Fig. 4A, yellow arrow) was lost in the PFHxS- and PFOS-treated oocytes. To examine the possibility of spindle migration failure in the PFAS-treated oocytes, we employed time-lapse confocal microscopy to track the spindle in live oocytes (Fig. 5). We observed partial spindle movement in PFHxS (30%) and PFOS-treated (22.22%) oocytes compared to control oocytes (4.76%). Importantly, we observed a significant increase in spindle migration failure in PFHxS (20%) and PFOS-treated oocytes (77.78%) when compared to untreated control oocytes (0%). Interestingly, we also found that spindle length-to-width ratio (Fig. 4C) and spindle-length-to-oocyte-diameter ratio (Fig. 4D) significantly increase in PFHxS- and PFOS-treated oocytes when compared to untreated controls. This spindle elongation potentially overcomes spindle migration failure. In addition, the chromosome misalignment rate was also increased dramatically as indicated by increased metaphase-plate width after PFHxS and PFOS exposure (Fig. 4E).

**Figure 4.**
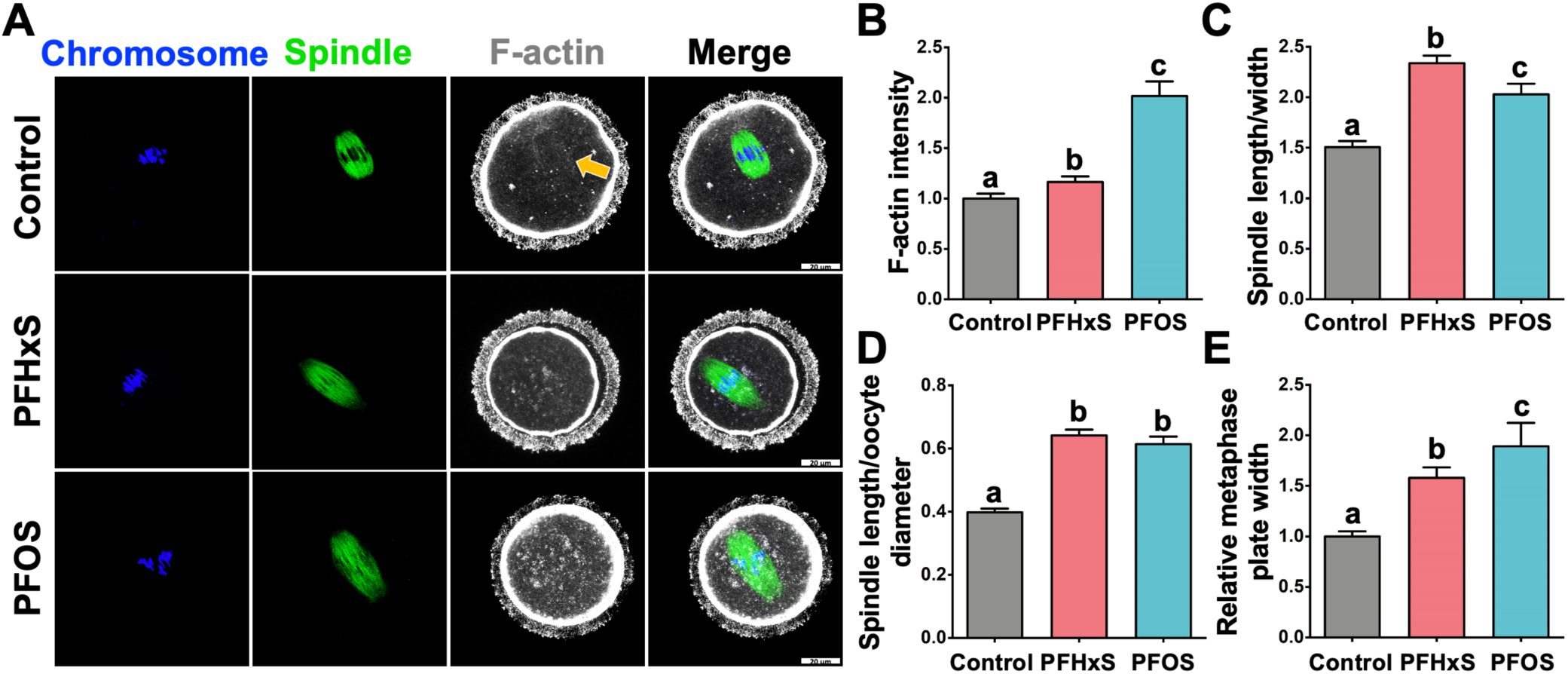
PFHxS and PFOS (600 µM) exposure causes misaligned chromosomes, elongated spindles, and disrupt F-actin organization. (A) Representative images of oocyte cytoskeleton structures in the control, 600 µM PFHxS-, and 600 µM PFOS-treated groups. Blue: DNA/chromosome; Green: microtubules (the spindle); White: F-actin. (B) F-actin intensity around the spindle regions, control: n = 21; PFHxS: n = 40; PFOS: n = 20. (C) Spindle length-to-width ratio. (D) Spindle-length-to-oocyte-diameter ratio. (E) Relative metaphase plate width. A total of 20 oocytes in the control, 35 oocytes in the PFHxS group, and 16 oocytes in the PFOS group were measured for (C), (D), and (E). Groups with different letters in the graphs (B-E) are significantly different (p < 0.05). Scale bar: 20 µm.

**Figure 5.**
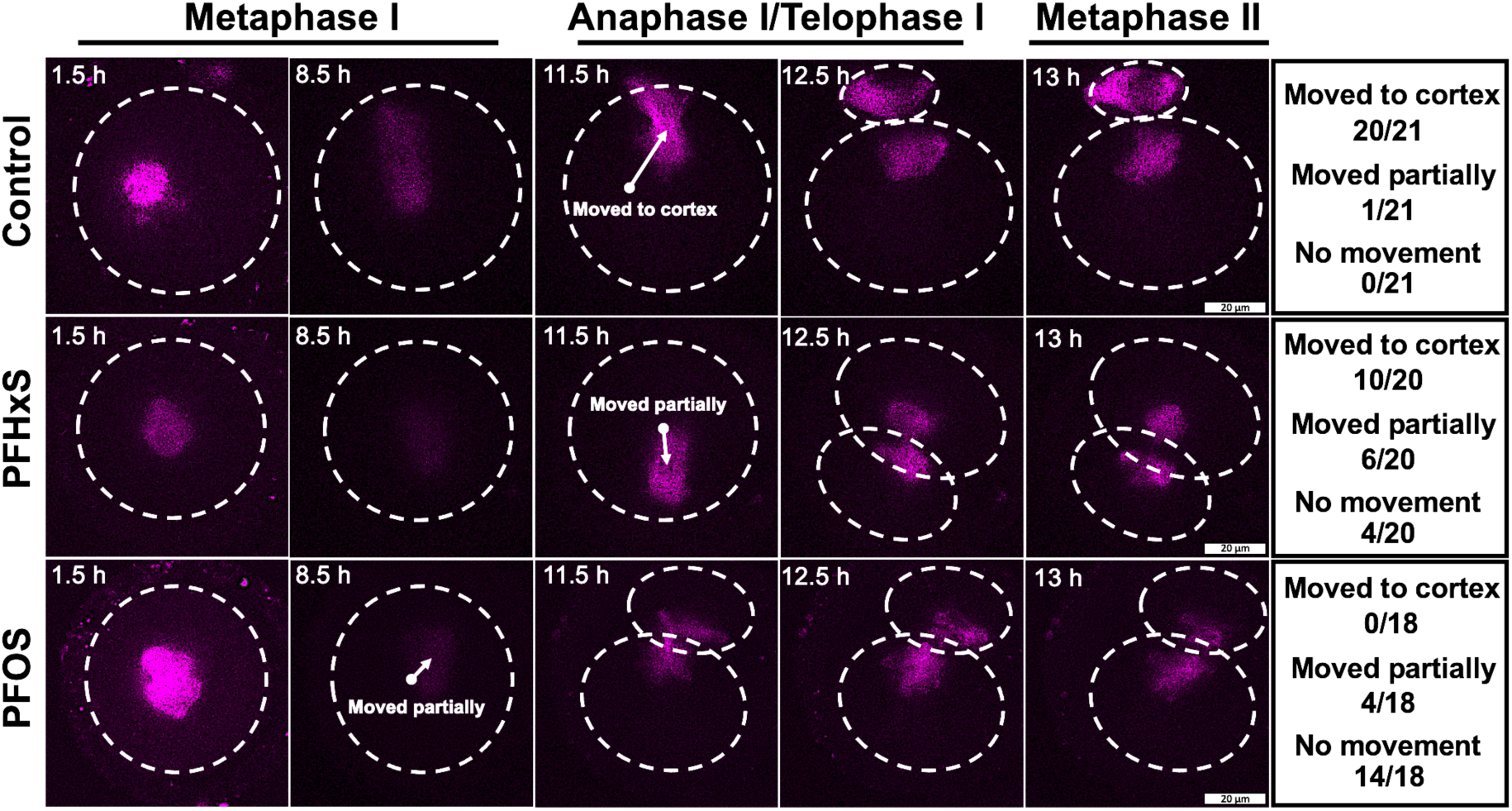
600 µM PFHxS and PFOS hindered peripheral spindle migration. Z-projection of time-lapse confocal imaging of oocytes cultured in the DMSO (control), 600 µM PFHxS-, and 600 µM PFOS-treated groups with SiR-tubulin (to label the spindle, magenta). Time-lapse imaging started at germinal vesicle breakdown (GVBD). White arrows indicate the movement of spindle. Table on the right summarizes the number of oocytes in each group that completely moved to the cortex region; partially moved to the cortex region; and had no movement. Scale bar: 20 µm.

## 4. Discussion

As persistent organic chemicals, PFAS have been detected ubiquitously, even in the liver samples from polar bears in Alaska.^50^ In humans, epidemiological studies show that human exposure to PFAS either prenatally or postnatally is related to reproductive defects in both males and females.^51–53^ Previous studies mostly focused on the toxicity of individual long-chain PFAS, such as PFOA and PFOS. Short chain PFAS such as PFBA were often neglected due to their relatively low bioaccumulation effect.^54^ In this study, using a mouse oocyte *in vitro* maturation system, we demonstrated that the toxicity of PFAS increases with the elongated carbon chain and the inclusion of a sulfonate group. Taking advantage of time-lapse confocal microscopy and immunocytochemistry, we also found aberrant cytoskeleton structure organization in PFAS-treated oocytes.

In terms of the carbon-chain-length effect, consistent with our finding, in other cell types including human colon carcinoma (HCT116)^55^ and human hepatocarcinoma (HepG2)^56^, the cytotoxicity of PFAS was positively correlated with the carbon-chain length when the length is smaller than 10. However, a weaker toxic effect was observed when the chain length further increased. We speculated that such a reversed-U-shape toxic effect derived from two aspects of PFAS’ characteristics. On one hand, PFAS are amphiphilic substances that can disturb cell membrane structures.^57^ Short-chain PFAS are less lipophilic than long-chain PFAS.^58^ Therefore, the “detergent effect” and bioaccumulation effect are less pronounced for short-chain PFAS than long-chain PFAS, which limit their toxicity. On the other hand, the capacity of PFAS to bind some receptors such as human PPARα ligand-binding domain will be reduced if the molecular size is too large^59^, restricting the toxicity of long-chain PFAS. Due to these two reasons, PFAS with middle length (around 10) are most toxic (Abstract Graphic). In the current study, we found an increasing toxicity for PFSA with carbon-chain length from 4 to 8 (solid curve in Abstract Graphic). However, whether the reversed-U-shape toxic effect exists in oocytes needs further investigation.

We also observed that PFSA toxicity is higher than that of PFCA, which is the case for other cell types including Sertoli cells.^60^ The higher cytotoxicity of PFSA is probably due to several reasons. First, the lipophilicity (octanol-water partition coefficient, KOW) of PFCA and PFSA are both very high, implying their ability to cross the cell membranes is limited by their transfer at a membrane/water interface (sorption affinity to artificial phospholipid membranes, KMW)^58^, which is higher in PFSA.^61^ To prove this, a novel method needs to be developed to precisely detect the amount of PFAS inside a single oocyte. Next, based on our observation under a time-lapse confocal microscopy: symmetrical oocyte division only occurred in the PFSA-treated groups. This indicates that PFSA hinders spindle migration. Furthermore, a sulfonate buffer can induce tubulin polymerization,^62^ leading to abnormal spindle morphology (Figure 5). In addition, both PFCA and PFSA have negatively charged heads and have the potential to bind to positively charged proteins like histone through electrostatic attraction. This effect is stronger for PFSA due to their lower pKa values (pKa of PFSA < < 0) than PFCA (pKa of PFCA < 4).^63^ Taken together, we concluded that PFSA are more toxic than PFCA with the same carbon-chain length.

We next selected 600 µM PFHxS and PFOS to study their toxic mechanisms. First, we observed mitochondrial dysfunction including an increased intracellular ROS level and diminishing MMP in PFHxS- and PFOS-treated oocytes, which have been reported for other types of PFAS like PFNA and PFOA, both in vivo^19–21^ and in vitro^64^. For PFOS exposure, Wei et al. also measured the expression of antioxidant enzyme levels inside oocytes. They found decreased expression of glutathione peroxidase (GSH-Px) and superoxide dismutase (SOD) but dramatically increased expression of catalase (CAT)^19^ in response to PFOS compared to control. PFHxS and PFOS can inhibit several cytochrome P450 enzyme (CYP) activities.^65^ This may explain the elevated ROS level observed in response to the chemicals in our study. We propose that PFHxS and PFOS can cause the uncoupling of the enzymatic cycle and excess ROS release (for more explanation, readers are referred to Veith and Moorthy, 2018)^66^. Similar pathways have been reported to induce oxidative stress like polychlorinated biphenyl (PCB).^67,68^ Excess ROS can cause many problems in oocytes, especially after GVBD, including DNA damage^17,20^ and early apoptosis (Supplemental Fig. 2). Another mitochondria-related problem is the decreased MMP level. MMP is important for ATP production^69^ and oocyte maturation. The decreased MMP (depolarization) after PFHxS and PFOS exposure implies damage to the mitochondrial structure. As surfactants, PFHxS and PFOS can disturb the mitochondrial membrane structures, increasing proton leakage of mitochondrial inner membrane.^70,71^ Such pathogenic proton leakage causes more protons to bypass ATP synthase, producing heat instead of ATP and decreasing MMP.^72^ Some caspase activators including cytochrome c, hsp 10, and hsp 60 can also be released through damaged mitochondrial membrane,^73,74^ which can activate executionary caspases leading to early apoptosis.

Spindle assembly and migration are also important for successful oocyte maturation and fertilization. In normally developed metaphase I oocytes, microtubules build up barrel-shaped spindles that facilitate chromosome alignment at the metaphase plate. Under the forces of F-actin, the spindles migrate from the cell center to a sub-cortical location to allow an asymmetric division.^75^ However, we found many oocytes in PFOS-treated oocytes underwent symmetric division, instead. Our data showed significant increases of F-actin fluorescent intensity around spindle regions after PFSA treatment, together with the loss of key F-actin structures like spindle-associated actin cage. Therefore, we propose that PFSA might act as an F-actin-stabilizing agent like jasplakinolide, a cyclo-depsipeptide actin stabilizer.^76^ Indeed, treating mouse oocytes with jasplakinolide prevented spindle migration during meiosis I.^46^ Therefore, these thick non-dynamic actin filaments could block the migration of the spindle leading to large PB extrusion in PFSA-treated oocytes.^19^ Another unique phenotype we observed in the PFSA treatment groups was elongated spindles. This could be explained by the fact that sulfonate buffers can induce the polymerization of tubulin to enlarge spindles.^61^ However, Verlhac et al. proposed that the elongated spindle could be a compensation for oocytes with unmigrated spindles to extrude a normal-sized PB.^77^ For example, they found in *mos*^*-/-*^ oocytes which lack mitogen-activated protein (MAP) kinase activity^78,79^, non-migrating spindles elongates so that one pole can be closer to the cortex while the other pole remained near the oocyte center.^77^ Therefore, some of the *mos*^*-/-*^ oocytes still can extrude their first PBs of normal sizes, which is also the case for PFHxS- and PFOS-treated oocytes.

In summary, from a female reproduction perspective, we demonstrated that PFAS with a longer chain and a sulfonate group are more toxic and revealed their toxic mechanisms. From an environmental health perspective, short chain PFAS may be less toxic than long chain PFAS according to our results. However, the health concerns regarding their endocrine-disrupting effects still need more research.^80^

## ASSOCIATED CONTENT

### Supporting Information

PFBA, PFHxA, and PFOA at 600 µM showed no effects on mouse oocyte *in vitro* maturation (Figure S1). Time-lapse microscopy showed oocyte maturation process in the control, 600 µM PFHxS, and 600 µM PFOS groups (Figure S2).

## AUTHOR INFORMATION

### Notes

The authors declare no competing financial interest.

## ACKNOWLEDGMENT

We would like to thank Dr. Joseph Irudayaraj for kindly sharing all PFAS chemicals used in this study. We also thank Drs. Michael Spinella, Sarah Freemantle, and Wenyan Mei for their technical support and Dr. Jodi Flaws for her comments on this manuscript. This work was supported by National Institutions of Health (NIH) R00HD082375, R01GM135549 and R35GM142537. JF acknowledges the IETP Toxicology Scholar Award.

## Abstract Graphic

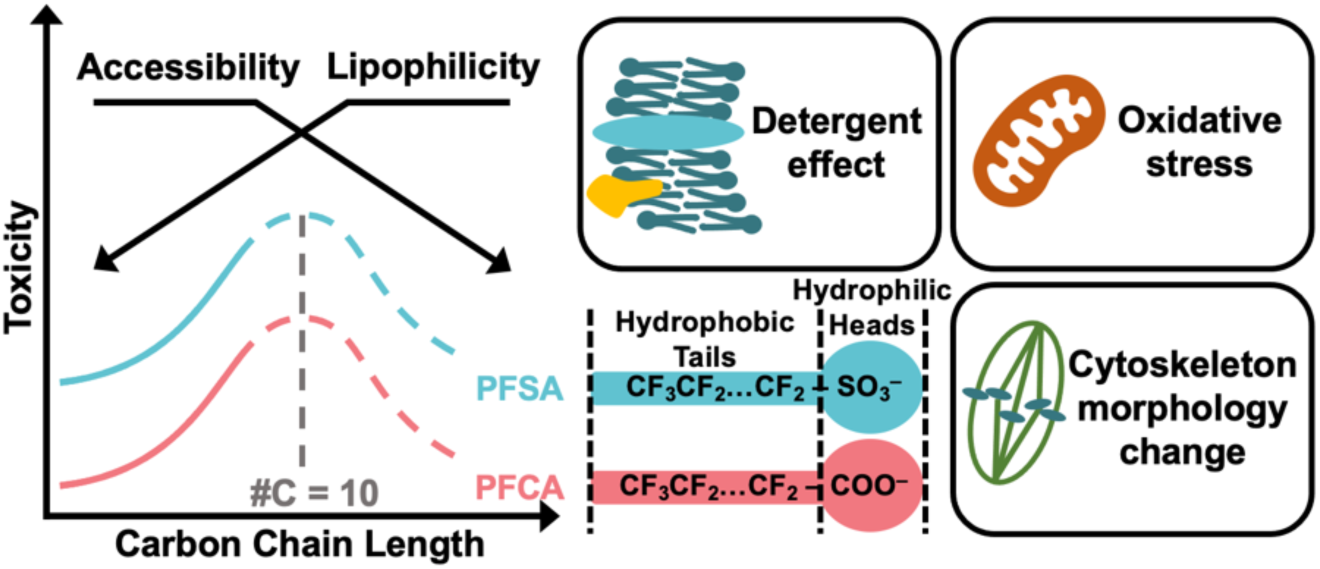

## REFERENCES

(1) Buck, R. C.; Franklin, J.; Berger, U.; Conder, J. M.; Cousins, I. T.; Voogt, P. de; Jensen, A. A.; Kannan, K.; Mabury, S. A.; van Leeuwen, S. P. J. Perfluoroalkyl and Polyfluoroalkyl Substances in the Environment: Terminology, Classification, and Origins. Integrated Environmental Assessment and Management 2011, 7 (4), 513–541.

(2) Glüge, J.; Scheringer, M.; Cousins, I. T.; Dewitt, J. C.; Goldenman, G.; Herzke, D.; Lohmann, R.; Ng, C. A.; Trier, X.; Wang, Z. An Overview of the Uses of Per- and Polyfluoroalkyl Substances (PFAS). Environmental Science: Processes & Impacts 2020, 22 (12), 2345–2373.

(3) Cousins, I. T.; Dewitt, J. C.; Glüge, J.; Goldenman, G.; Herzke, D.; Lohmann, R.; Ng, C. A.; Scheringer, M.; Wang, Z. The High Persistence of PFAS Is Sufficient for Their Management as a Chemical Class. Environ Sci Process Impacts 2020, 22 (12), 2307–2312.

(4) Haukås, M.; Berger, U.; Hop, H.; Gulliksen, B.; Gabrielsen, G. W. Bioaccumulation of Per- and Polyfluorinated Alkyl Substances (PFAS) in Selected Species from the Barents Sea Food Web. Environmental Pollution 2007, 148 (1), 360–371.

(5) Poothong, S.; Papadopoulou, E.; Padilla-Sánchez, J. A.; Thomsen, C.; Haug, L. S. Multiple Pathways of Human Exposure to Poly- and Perfluoroalkyl Substances (PFASs): From External Exposure to Human Blood. Environment International 2020, 134, 105244.

(6) Foguth, R.; Sepúlveda, M. S.; Cannon, J. Per- and Polyfluoroalkyl Substances (PFAS) Neurotoxicity in Sentinel and Non-Traditional Laboratory Model Systems: Potential Utility in Predicting Adverse Outcomes in Human Health. Toxics 2020, 8 (2), 42.

(7) Eggers Pedersen, K.; Basu, N.; Letcher, R.; Greaves, A. K.; Sonne, C.; Dietz, R.; Styrishave, B. Brain Region-Specific Perfluoroalkylated Sulfonate (PFSA) and Carboxylic Acid (PFCA) Accumulation and Neurochemical Biomarker Responses in East Greenland Polar Bears (Ursus Maritimus). Environ Res 2015, 138, 22–31.

(8) Gaballah, S.; Swank, A.; Sobus, J. R.; Howey, X. M.; Schmid, J.; Catron, T.; McCord, J.; Hines, E.; Strynar, M.; Tal, T. Evaluation of Developmental Toxicity, Developmental Neurotoxicity, and Tissue Dose in Zebrafish Exposed to GenX and Other PFAS. Environmental Health Perspectives 2020, 128 (4).

(9) DeWitt, J. C.; Blossom, S. J.; Schaider, L. A. Exposure to Per-Fluoroalkyl and Polyfluoroalkyl Substances Leads to Immunotoxicity: Epidemiological and Toxicological Evidence. Journal of Exposure Science & Environmental Epidemiology 2018, 29 (2), 148–156.

(10) Jin, R.; McConnell, R.; Catherine, C.; Xu, S.; Walker, D. I.; Stratakis, N.; Jones, D. P.; Miller, G. W.; Peng, C.; Conti, D. v.; Vos, M. B.; Chatzi, L. Perfluoroalkyl Substances and Severity of Nonalcoholic Fatty Liver in Children: An Untargeted Metabolomics Approach. Environ Int 2020, 134, 105220.

(11) Tarapore, P.; Ouyang, B. Perfluoroalkyl Chemicals and Male Reproductive Health: Do PFOA and PFOS Increase Risk for Male Infertility? International Journal of Environmental Research and Public Health 2021, 18 (7), 3794.

(12) Chambers, W. S.; Hopkins, J. G.; Richards, S. M. A Review of Per- and Polyfluorinated Alkyl Substance Impairment of Reproduction. Frontiers in Toxicology 2021, 3, 53.

(13) Kato, K.; Wong, L. Y.; Chen, A.; Dunbar, C.; Webster, G. M.; Lanphear, B. P.; Calafat, A. M. Changes in Serum Concentrations of Maternal Poly- and Perfluoroalkyl Substances over the Course of Pregnancy and Predictors of Exposure in a Multiethnic Cohort of Cincinnati, Ohio Pregnant Women during 2003-2006. Environmental Science and Technology 2014, 48 (16), 9600–9608.

(14) Ding, N.; Harlow, S. D.; Randolph, J. F.; Loch-Caruso, R.; Park, S. K. Perfluoroalkyl and Polyfluoroalkyl Substances (PFAS) and Their Effects on the Ovary. Human Reproduction Update 2020, 26 (5), 724–752.

(15) He, M.; Zhang, T.; Yang, Y.; Wang, C. Mechanisms of Oocyte Maturation and Related Epigenetic Regulation. Frontiers in Cell and Developmental Biology 2021, 9, 654028.

(16) Jiao, X.; Gonsioroski, A.; Flaws, J. A.; Qiao, H. Iodoacetic Acid Disrupts Mouse Oocyte Maturation by Inducing Oxidative Stress and Spindle Abnormalities. Environmental Pollution 2021, 268, 115601.

(17) Jo, Y. J.; Yoon, S. bin; Park, B. J.; Lee, S. il; Kim, K. J.; Kim, S. Y.; Kim, M.; Lee, J. K.; Lee, S. Y.; Lee, D. H.; Kwon, T.; Son, Y.; Lee, J. R.; Kwon, J.; Kim, J. S. Particulate Matter Exposure During Oocyte Maturation: Cell Cycle Arrest, ROS Generation, and Early Apoptosis in Mice. Frontiers in Cell and Developmental Biology 2020, 8, 1402.

(18) López-Arellano, P.; López-Arellano, K.; Luna, J.; Flores, D.; Jiménez-Salazar, J.; Gavia, G.; Teteltitla, M.; Rodríguez, J. J.; Domínguez, A.; Casas, E.; Bahena, I.; Betancourt, M.; González, C.; Ducolomb, Y.; Bonilla, E. Perfluorooctanoic Acid Disrupts Gap Junction Intercellular Communication and Induces Reactive Oxygen Species Formation and Apoptosis in Mouse Ovaries. Environmental Toxicology 2019, 34 (1), 92–98.

(19) Wei, K. N.; Wang, X. J.; Zeng, Z. C.; Gu, R. T.; Deng, S. Z.; Jiang, J.; Xu, C. L.; Li, W.; Wang, H. L. Perfluorooctane Sulfonate Affects Mouse Oocyte Maturation in Vitro by Promoting Oxidative Stress and Apoptosis Induced Bymitochondrial Dysfunction. Ecotoxicology and Environmental Safety 2021, 225, 112807.

(20) Jiao, X.; Liu, N.; Xu, Y.; Qiao, H. Perfluorononanoic Acid Impedes Mouse Oocyte Maturation by Inducing Mitochondrial Dysfunction and Oxidative Stress. Reproductive Toxicology 2021, 104, 58–67.

(21) Guo, C.; Zhao, Z.; Zhao, K.; Huang, J.; Ding, L.; Huang, X.; Meng, L.; Li, L.; Wei, H.; Zhang, S. Perfluorooctanoic Acid Inhibits the Maturation Rate of Mouse Oocytes Cultured in Vitro by Triggering Mitochondrial and DNA Damage. Birth Defects Research 2021, 113 (14), 1074–1083.

(22) Sankar, A.; Lerdrup, M.; Manaf, A. KDM4A Regulates the Maternal-to-Zygotic Transition by Protecting Broad H3K4me3 Domains from H3K9me3 Invasion in Oocytes. Nature Cell Biology 2020, 22, 380–388.

(23) Arne Dahl, J.; Jung, I.; Aanes, H.; Greggains, G. D.; manaf, A.; Lerdrup, mads; Li, G.; Kuan, samantha; Li, bin; Young Lee, A.; preissl, sebastian; Jermstad, I.; Haugland Haugen, mads; suganthan, R.; bjørås, magnar; Hansen, K.; Tomas Dalen, K., Fedorcsak, peter; Ren, bing; Klungland, A. Broad Histone H3K4me3 Domains in Mouse Oocytes Modulate Maternal-to-Zygotic Transition. Nature 2016, 537, 548–552.

(24) Kirillova, A.; Smitz, J. E. J.; Sukhikh, G. T.; Mazunin, I. The Role of Mitochondria in Oocyte Maturation. Cells 2021, 10 (9), 2484.

(25) Wang, L.-Q.; Liu, T.; Yang, S.; Sun, L.; Zhao, Z.-Y.; Li, L.-Y.; She, Y.-C.; Zheng, Y.-Y.; Ye, X.-Y.; Bao, Q.; Dong, G.-H.; Li, C.-W.; Cui, J. Perfluoroalkyl Substance Pollutants Activate the Innate Immune System through the AIM2 Inflammasome. Nature Communications 2021, 12, 2915

(26) Choi, E. M.; Suh, K. S.; Rhee, S. Y.; Oh, S.; Woo, J. T.; Kim, S. W.; Kim, Y. S.; Pak, Y. K.; Chon, S. Perfluorooctanoic Acid Induces Mitochondrial Dysfunction in MC3T3-E1 Osteoblast Cells. Journal of Environmental Science and Health 2016, 52 (3), 281–289.

(27) Kleszczyński, K.; Stepnowski, P. Mechanism of Cytotoxic Action of Perfluorinated Acids. II. Disruption of Mitochondrial Bioenergetics Pharmaceuticals and Their Transformation Products in the Environment: Analytics, Ecotoxicology and Risk Assessment View Project PASSIL-an Innovative Passive Sampling Technique Using Ionic Liquids View Project. Toxicology and Applied Pharmacology 2009. 235, 182–190

(28) Hale, B. J.; Li, Y.; Adur, M. K.; Ross, J. W. Inhibition of Germinal Vesicle Breakdown Using IBMX Increases MicroRNA-21 in the Porcine Oocyte. Reproductive Biology and Endocrinology 2020, 18 (1), 39.

(29) Schindelin, J.; Arganda-Carreras, I.; Frise, E.; Kaynig, V.; Longair, M.; Pietzsch, T.; Preibisch, S.; Rueden, C.; Saalfeld, S.; Schmid, B.; Tinevez, J. Y.; White, D. J.; Hartenstein, V.; Eliceiri, K.; Tomancak, P.; Cardona, A. Fiji: An Open-Source Platform for Biological-Image Analysis. Nature Methods 2012 9 (7), 676–682.

(30) Londoño-Vásquez, D.; Rodriguez-Lukey, K.; Behura, S. K.; Balboula, A. Z. Microtubule Organizing Centers Regulate Spindle Positioning in Mouse Oocytes. Developmental Cell 2022, 57 (2), 197–211.

(31) Balboula, A. Z.; Nguyen, A. L.; Gentilello, A. S.; Quartuccio, S. M.; Drutovic, D.; Solc, P.; Schindler, k. Haspin kinase regulates microtubule-organizing center clustering and stability through Aurora kinase C in mouse oocytes. J Cell Sci 2016, 129 (19): 3648–3660.

(32) Zhou, D.; Shen, X.; Gu, Y.; Zhang, N.; Li, T.; Wu, X.; Lei, L. Effects of Dimethyl Sulfoxide on Asymmetric Division and Cytokinesis in Mouse Oocytes. BMC Developmental Biology 2014, 14, 1–7.

(33) Jo, Y. J.; Yoon, S. bin; Park, B. J.; Lee, S. il; Kim, K. J.; Kim, S. Y.; Kim, M.; Lee, J. K.; Lee, S. Y.; Lee, D. H.; Kwon, T.; Son, Y.; Lee, J. R.; Kwon, J.; Kim, J. S. Particulate Matter Exposure During Oocyte Maturation: Cell Cycle Arrest, ROS Generation, and Early Apoptosis in Mice. Frontiers in Cell and Developmental Biology 2020, 8, 1402.

(34) Langenbach, B.; Wilson, M. Per- and Polyfluoroalkyl Substances (PFAS): Significance and Considerations within the Regulatory Framework of the USA. International Journal of Environmental Research and Public Health 2021, 18 (21), 11142.

(35) Morrell, C. N. Reactive Oxygen Species: Finding the Right Balance. Circ Res 2008, 103 (6), 571.

(36) Das, K.; Roychoudhury, A. Reactive Oxygen Species (ROS) and Response of Antioxidants as ROS-Scavengers during Environmental Stress in Plants. Frontiers in Environmental Science 2014, 2, 53.

(37) Kalyanaraman, B.; Darley-Usmar, V.; Davies, K. J. A.; Dennery, P. A.; Forman, H. J.; Grisham, M. B.; Mann, G. E.; Moore, K.; Roberts, L. J.; Ischiropoulos, H. Measuring Reactive Oxygen and Nitrogen Species with Fluorescent Probes: Challenges and Limitations. Free Radic Biol Med 2012, 52 (1), 1–6.

(38) Tetz, L. M.; Kamau, P. W.; Cheng, A. A.; Meeker, J. D.; Loch-Caruso, R. Troubleshooting the Dichlorofluorescein Assay to Avoid Artifacts in Measurement of Toxicant-Stimulated Cellular Production of Reactive Oxidant Species. J Pharmacol Toxicol Methods 2013, 67 (2), 56.

(39) Suzuki-Karasaki, M.; Ochiai, T.; Suzuki-Karasaki, Y. Crosstalk between Mitochondrial ROS and Depolarization in the Potentiation of TRAIL-Induced Apoptosis in Human Tumor Cells. International Journal of Oncology 2014, 44 (2), 616–628.

(40) Matsuyama, S.; Reed, J. C. Mitochondria-Dependent Apoptosis and Cellular PH Regulation. Cell Death & Differentiation 2000 7 (12), 1155–1165.

(41) Al-Zubaidi, U.; Liu, J.; Cinar, O.; Robker, R. L.; Adhikari, D.; Carroll, J. The Spatio-Temporal Dynamics of Mitochondrial Membrane Potential during Oocyte Maturation. Molecular Human Reproduction 2019, 25 (11), 695.

(42) Palmeira, C. M.; Ramalho-Santos, J. Mitochondrial Dysfunction in Reproductive and Developmental Toxicity. Reproductive and Developmental Toxicology 2017, 1023–1035.

(43) Ebner, T.; Yaman, C.; Moser, M.; Sommergruber, M.; Feichtinger, O.; Tews, G. Prognostic Value of First Polar Body Morphology on Fertilization Rate and Embryo Quality in Intracytoplasmic Sperm Injection. Human Reproduction 2000, 15 (2), 427–430.

(44) Younis, J. S.; Radin, O.; Izhaki, I.; Ben-Ami, M. Does First Polar Body Morphology Predict Oocyte Performance during ICSI Treatment? Journal of Assisted Reproduction and Genetics 2009, 26 (11–12), 561.

(45) Rose, B. I.; Laky, D. Polar Body Fragmentation in IVM Oocytes Is Associated with Impaired Fertilization and Embryo Development. Journal of Assisted Reproduction and Genetics 2013, 30 (5), 679.

(46) Terada, Y.; Simerly, C. and Schatten, G. Microfilament Stabilization by Jasplakinolide Arrests Oocyte Maturation, Cortical Granule Exocytosis, Sperm Incorporation Cone Resorption, and Cell-Cycle Progression, but not DNA Replication, during Fertilization in Mice. Mol Reprod Dev 2000, 56, 89–98.

(47) Azoury, J.; Lee, K.W.; Georget, V.; Rassinier, P.; Leader, B.; Verlhac, M.-H. Spindle Positioning in Mouse Oocytes Relies on a Dynamic Meshwork of Actin Filaments. Current Biology 2008, 18 (19), 1514–1519.

(48) Schuh, M.; Ellenberg, J. A New Model for Asymmetric Spindle Positioning in Mouse Oocytes. Current Biology 2008, 18 (24), 1986–1992.

(49) Longo, F. J.; Chen, D.-Y. Development of Cortical Polarity in Mouse Eggs: Involvement of the Meiotic Apparatus. Developmental Biology 1985, 107 (2), 382–394.

(50) Giesy, J. P.; Kannan, K. Global Distribution of Perfluorooctane Sulfonate in Wildlife. Environmental Science and Technology 2001, 35 (7), 1339–1342.

(51) Eick, S. M.; Hom Thepaksorn, E. K.; Izano, M. A.; Cushing, L. J.; Wang, Y.; Smith, S. C.; Gao, S.; Park, J. S.; Padula, A. M.; Demicco, E.; Valeri, L.; Woodruff, T. J.; Morello-Frosch, R. Associations between Prenatal Maternal Exposure to Per- And Polyfluoroalkyl Substances (PFAS) and Polybrominated Diphenyl Ethers (PBDEs) and Birth Outcomes among Pregnant Women in San Francisco. Environmental Health: A Global Access Science Source 2020, 19 (1), 1–12.

(52) Petersen, K. U.; Larsen, J. R.; Deen, L.; Flachs, E. M.; Hærvig, K. K.; Hull, S. D.; Bonde, J. P. E.; Tøttenborg, S. S. Per- and Polyfluoroalkyl Substances and Male Reproductive Health: A Systematic Review of the Epidemiological Evidence. Journal of Toxicology and Environmental Health -Part B: Critical Reviews 2020, 23 (6), 276–291.

(53) Rickard, B. P.; Rizvi, I.; Fenton, S. E. Per- and Poly-Fluoroalkyl Substances (PFAS) and Female Reproductive Outcomes: PFAS Elimination, Endocrine-Mediated Effects, and Disease. Toxicology 2022, 465, 153031.

(54) Abraham, K.; El-Khatib, A. H.; Schwerdtle, T.; Monien, B. H. Perfluorobutanoic Acid (PFBA): No High-Level Accumulation in Human Lung and Kidney Tissue. International Journal of Hygiene and Environmental Health 2021, 237, 113830.

(55) Kleszczyński, K.; Gardzielewski, P.; Mulkiewicz, E.; Stepnowski, P.; Składanowski, A. C. Analysis of Structure–Cytotoxicity in Vitro Relationship (SAR) for Perfluorinated Carboxylic Acids. Toxicology in Vitro 2007, 21 (6), 1206–1211.

(56) Buhrke, T.; Kibellus, A.; Lampen, A. In Vitro Toxicological Characterization of Perfluorinated Carboxylic Acids with Different Carbon Chain Lengths. Toxicology Letters 2013, 218 (2), 97–104.

(57) Oldham, E. D.; Xie, W.; Farnoud, A. M.; Fiegel, J.; Lehmler, H. J. Disruption of Phosphatidylcholine Monolayers and Bilayers by Perfluorobutane Sulfonate. Journal of Physical Chemistry B 2012, 116 (33), 9999–10007.

(58) Jing, P.; Rodgers, P. J.; Amemiya, S. High Lipophilicity of Perfluoroalkyl Carboxylate and Sulfonate: Implications for Their Membrane Permeability. J Am Chem Soc 2009, 131 (6), 2290–2296.

(59) Wolf, C. J.; Takacs, M. L.; Schmid, J. E.; Lau, C.; Abbott, B. D. Activation of Mouse and Human Peroxisome Proliferator−Activated Receptor Alpha by Perfluoroalkyl Acids of Different Functional Groups and Chain Lengths. Toxicological Sciences 2008, 106 (1), 162–171.

(60) Wan, H. T.; Lai, K. P.; Wong, C. K. C. Comparative Analysis of PFOS and PFOA Toxicity on Sertoli Cells. Environmental Science and Technology 2020, 54 (6), 3465–3475.

(61) Droge, S. T. J. Membrane-Water Partition Coefficients to Aid Risk Assessment of Perfluoroalkyl Anions and Alkyl Sulfates. Environmental Science and Technology 2019, 53 (2), 760–770.

(62) Waxman, P. G.; del Campo, A. A.; Lowe, M. C.; Hamel, E. Induction of Polymerization of Purified Tubulin by Sulfonate Buffers. European Journal of Biochemistry 1981, 120 (1), 129–136.

(63) Rayne, S.; Forest, K. An Assessment of Organic Solvent Based Equilibrium Partitioning Methods for Predicting the Bioconcentration Behavior of Perfluorinated Sulfonic Acids, Carboxylic Acids, and Sulfonamides. Nature Proceedings, 2009.

(64) Chen, Y.; Zhou, L.; Xu, J.; Zhang, L.; Li, M.; Xie, X.; Xie, Y.; Luo, D.; Zhang, D.; Yu, X.; Yang, B.; Kuang, H. Maternal Exposure to Perfluorooctanoic Acid Inhibits Luteal Function via Oxidative Stress and Apoptosis in Pregnant Mice. Reproductive Toxicology 2017, 69, 159–166.

(65) Amstutz, V. H.; Cengo, A.; Sijm, D. T. H. M.; Vrolijk, M. F. The Impact of Legacy and Novel Perfluoroalkyl Substances on Human Cytochrome P450: An in Vitro Study on the Inhibitory Potential and Underlying Mechanisms. Toxicology 2022, 468, 153116.

(66) Veith, A.; Moorthy, B. Role of Cytochrome P450s in the Generation and Metabolism of Reactive Oxygen Species. Current Opinion in Toxicology 2018, 7, 44–51.

(67) Green, R. M.; Hodges, N. J.; Chipman, J. K.; O’Donovan, M. R.; Graham, M. Reactive Oxygen Species from the Uncoupling of Human Cytochrome P450 1B1 May Contribute to the Carcinogenicity of Dioxin-like Polychlorinated Biphenyls. Mutagenesis 2008, 23 (6), 457–463.

(68) Schlezinger, J. J.; Struntz, W. D. J.; Goldstone, J. v.; Stegeman, J. J. Uncoupling of Cytochrome P450 1A and Stimulation of Reactive Oxygen Species Production by Co-Planar Polychlorinated Biphenyl Congeners. Aquatic Toxicology 2006, 77 (4), 422–432.

(69) Zorova, L. D.; Popkov, V. A.; Plotnikov, E. Y.; Silachev, D. N.; Pevzner, I. B.; Jankauskas, S. S.; Babenko, V. A.; Zorov, S. D.; Balakireva, A. v.; Juhaszova, M.; Sollott, S. J.; Zorov, D. B. Mitochondrial Membrane Potential. Analytical Biochemistry 2018, 552, 50–59.

(70) Starkov, A. A.; Wallace, K. B. Structural Determinants of Fluorochemical-Induced Mitochondrial Dysfunction. Toxicological Sciences 2002, 66, 244–252.

(71) Suh, K. S.; Choi, E. M.; Kim, Y. J.; Hong, S. M.; Park, S. Y.; Rhee, S. Y.; Oh, S.; Kim, S. W.; Pak, Y. K.; Choe, W.; Chon, S. Perfluorooctanoic Acid Induces Oxidative Damage and Mitochondrial Dysfunction in Pancreatic β-Cells. Molecular Medicine Reports 2017, 15 (6), 3871–3878.

(72) Mailloux, R. J.; Harper, M. E. Uncoupling Proteins and the Control of Mitochondrial Reactive Oxygen Species Production. Free Radical Biology and Medicine 2011, 51 (6), 1106–1115.

(73) Ly, J. D.; Grubb, D. R.; Lawen, A. The Mitochondrial Membrane Potential (Δψm) in Apoptosis; an Update. Apoptosis 2003, 8 (2), 115–128.

(74) Kroemer, G.; Reed, J. C. Mitochondrial Control of Cell Death. Nature Medicine 2000, 6 (5), 513–519.

(75) Duan, X.; Li, Y.; Yi, K.; Guo, F.; Wang, H. Y.; Wu, P. H.; Yang, J.; Mair, D. B.; Morales, E. A.; Kalab, P.; Wirtz, D.; Sun, S. X.; Li, R. Dynamic Organelle Distribution Initiates Actin-Based Spindle Migration in Mouse Oocytes. Nature Communications 2020, 11 (1), 1–15.

(76) Holzinger, A. Jasplakinolide: An Actin-Specific Reagent That Promotes Actin Polymerization. Methods Mol Biol 2009, 586, 71–87.

(77) Verlhac, M. H.; Lefebvre, C.; Guillaud, P.; Rassinier, P.; Maro, B. Asymmetric Division in Mouse Oocytes: With or without Mos. Current Biology 2000, 10 (20), 1303–1306.

(78) Carlton, H.; Udy, M. B. L.; Evans, G. B.; Hashimoto, M. J.; Watanabe, N.; Furuta, N.; Tamemoto, Y.; Sagata, H.; Yokoyama, N.; Okazaki, M.; Nagayoshi, K.; Takeda, M.; Ikawa, N.; Aizawa, Y. The Mos/Mitogen-Activated Protein Kinase (MAPK) Pathway Regulates the Size and Degradation of the First Polar Body in Maturing Mouse Oocytes. Proc Natl Acad Sci USA 1996, 93 (14), 7032–7035.

(79) Verlhac, M. H.; Kubiak, J. Z.; Weber, M.; Géraud, G.; Colledge, W. H.; Evans, M. J.; Maro, B. Mos Is Required for MAP Kinase Activation and Is Involved in Microtubule Organization during Meiotic Maturation in the Mouse. Development 1996, 122 (3), 815–822.

(80) Nian, M.; Luo, K.; Luo, F.; Aimuzi, R.; Huo, X.; Chen, Q.; Tian, Y.; Zhang, J. Association between Prenatal Exposure to PFAS and Fetal Sex Hormones: Are the Short-Chain PFAS Safer? Environmental Science and Technology 2020, 54 (13), 8291–8299.

