## Supplemental Figures for "Adverse PFAS effects on mouse oocyte *in vitro* maturation are associated with carbon-chain length and inclusion of a sulfonate group"

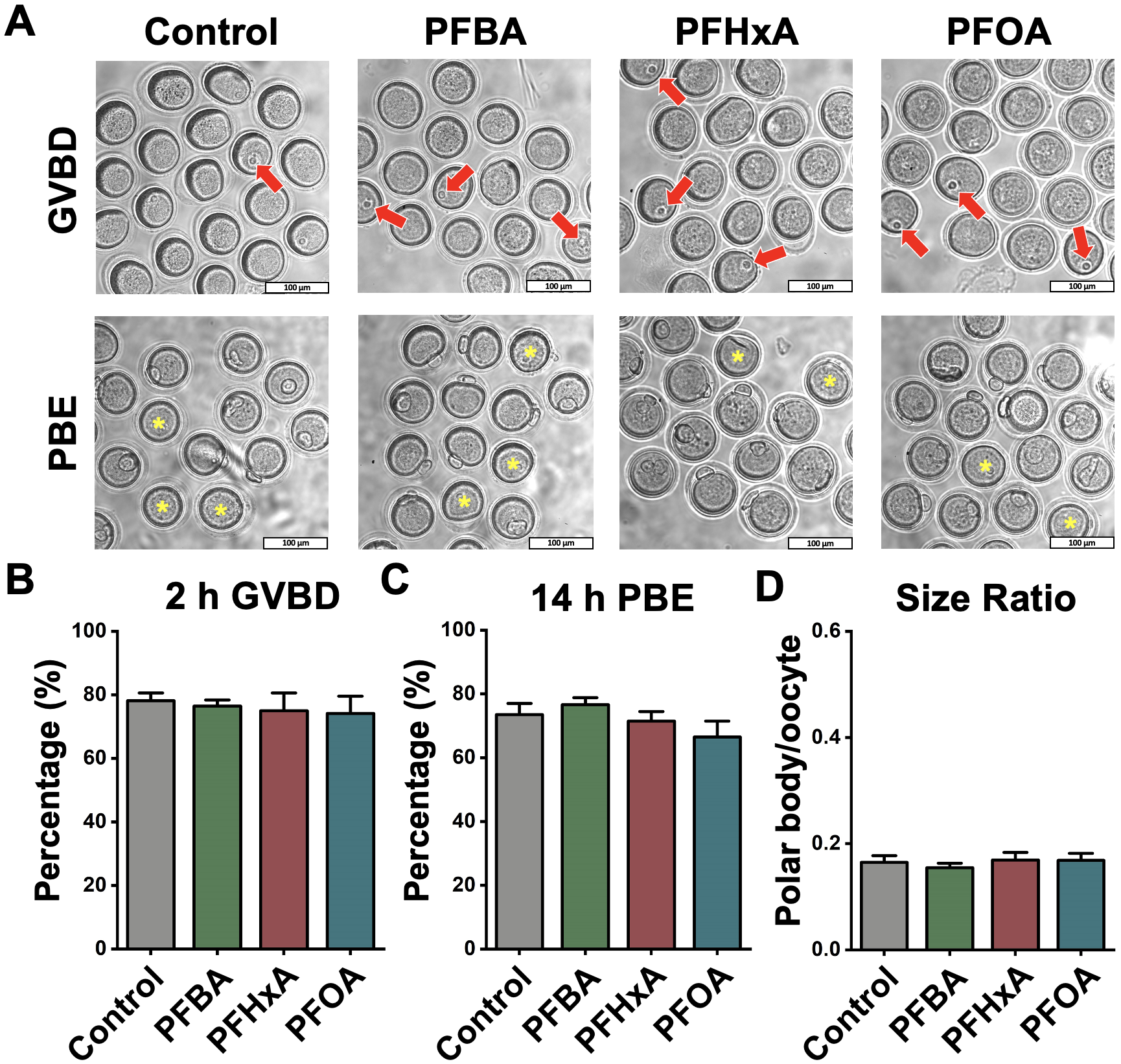

**Supplemental Figure 1.** PFCA at 600 µM showed no effects on mouse oocyte *in vitro* maturation. (A) Representative images show GVBD and PBE of four treatment groups (untreated, 600 μM PFBA, 600 μM PFHxA, and 600 μM PFOA). The red arrows indicate oocytes that retained their germinal vesicles after 2 hours of culture. The yellow asterisks indicate oocytes that did not extrude a PB after 14 hours of culture. Scale bar, 100 μm. (B) The rates of GVBD in the control and PFCA-treated groups. (C) The rates of PBE in the control and PFCA-treated groups. A total of 107 oocytes in the control group, 117 oocytes in the PFBA-treated group, 124 oocytes in the PFHxA-treated group, and 144 oocytes in the PFOA-treated group were analyzed to calculate the GVBD and PBE rates. (D) The size ratios of PBs to oocytes in the control and PFCA-treated groups. A total of 41 oocytes in the control group, 52 oocytes in the PFBA-treated group, 56 oocytes in the PFHxA-treated group, and 63 oocytes in the PFOA-treated group were measured to calculate the size ratios. Data in bar chart were presented as mean ± SEM. All groups had at least 3 independent groups.

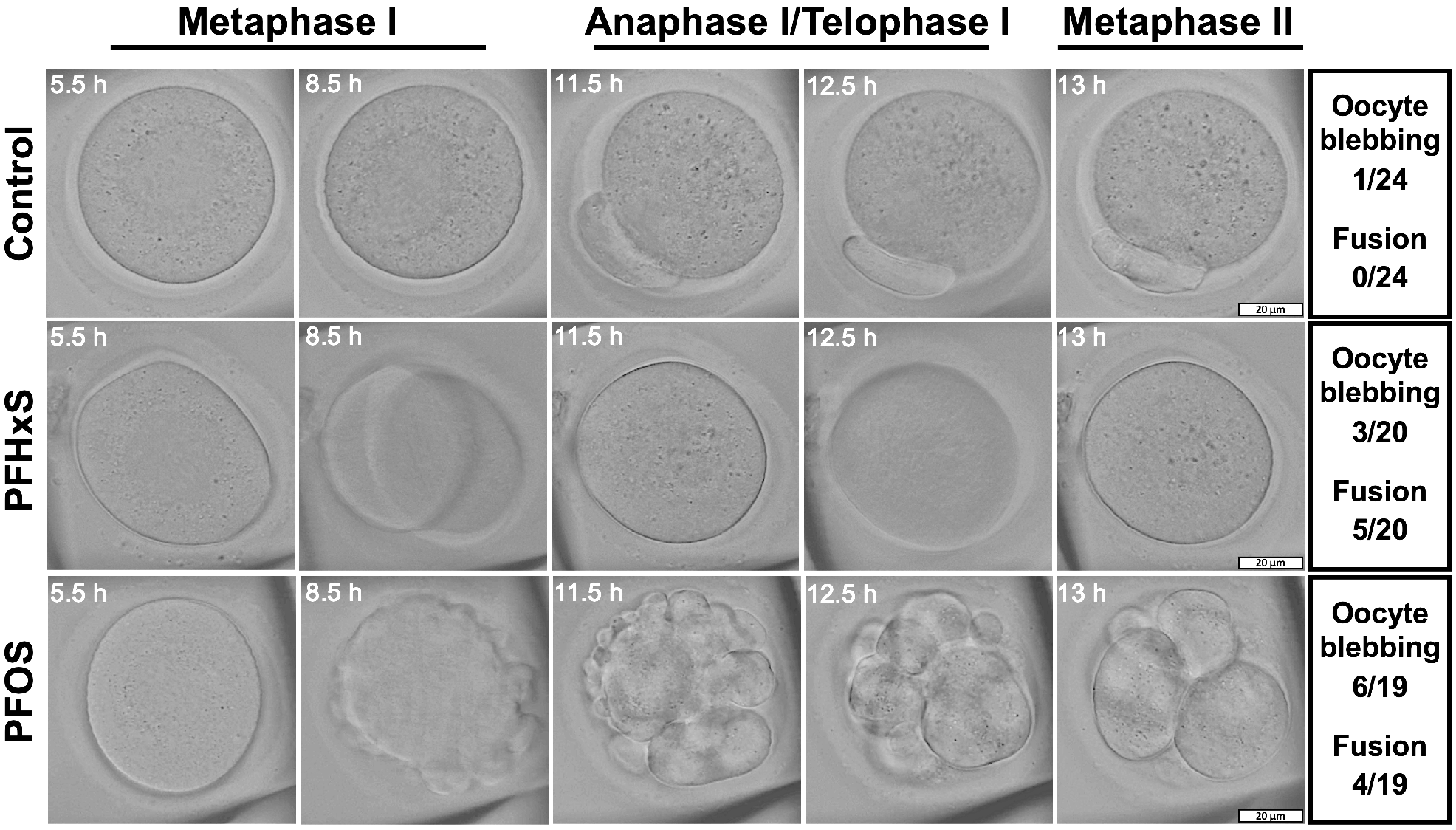

**Supplemental Figure 2.** Time-lapse bright-field images showing oocyte maturation process in the control, 600 µM PFHxS, and 600 µM PFOS groups. The oocyte in the control group extruded normal first polar body (PB) at anaphase I/telophase I. However, the oocyte in the PFHxS group extruded its large polar body prior to its resorption (oocyte and PB failure, cytokinesis failure). The oocyte in PFOS group underwent severe cytoplasmic blebbing followed cytokinesis defects. The table on the right listed the numbers of oocytes that experienced blebbing and fusion. Scale bar, 20 µm.
